# The fossil record of siliceous sponge spicules can be taken at face value

**DOI:** 10.64898/2026.01.27.702024

**Authors:** Sandy Y. Cui, Nicole S. Mizrahi, Shaily Rahman, Scott A. Nichols, Talia S. Karim, Carl Simpson

**Affiliations:** University of Colorado Museum of Natural History; Boulder, CO, USA 80309; Department of Geophysical Sciences, The University of Chicago; Chicago, IL, USA 60637; Department of Geological Sciences, University of Colorado Boulder; Boulder, CO, USA 80309; Institute of Arctic and Alpine Research (INSTAAR), University of Colorado Boulder; Boulder, CO, USA 80309; Department of Biological Sciences, University of Denver; Denver, CO 80208

## Abstract

Modern sponges (Porifera) diverged by the Cryogenian, but their silicious skeletons do not appear in the fossil record until one hundred million years later, a time-span termed the “spicule gap” and thought to be a taphonomic artifact even though sponges convergently evolved siliceous spicules. Due to sponges’ position in animal phylogeny and important role in regulating ocean chemistry, the timing of their biomineralization has major implications for the changing tempo and mode of Earth systems as animals radiate. In a comprehensive dataset of Ediacaran and Cambrian sponges, we find that spicules are readily preserved in Cambrian environments more extreme than those of the Ediacaran. Given the convergent evolution of siliceous spicules, we find that the fossil record accurately represents when spicules first evolved in the different sponge lineages.

## Introduction

Due to the early-diverging position of sponges within animal phylogeny, an understanding of when sponges, and in particular their siliceous spicules, evolved provides important insight into the tempo and mode of animal radiation and the role that skeletal origins played in the Cambrian Explosion. The inferred presence of sponges in the geological past, through biomarkers (1-5) and molecular phylogenetic divergence times (6-8), predates the appearance of body fossils and spicules in the rock record by well over 100 million years (9-11, Fig. 1). New genomic insights suggest that despite demosponges and hexactinellids being sister groups, their silicious spicules are convergent rather than homologous (12, 13). Despite this current understanding of the sponge phylogeny and the convergent evolution of siliceous spicules, we still do not know when spicules evolved. A face-value interpretation of the fossil record may nevertheless imply a substantial missing record of spicules (termed the “spicule gap”; 9), the length of which depends on when the sister groups demospongia and hexactinellida convergently evolved their siliceous spicules. The inferred divergence between demosponges and hexactinellids is likely to be deep, in the early Ediacaran or even Cryogenian (6, 8, 14, 15), but it remains an open question when spicules evolved in each lineage.

**Figure 1:**
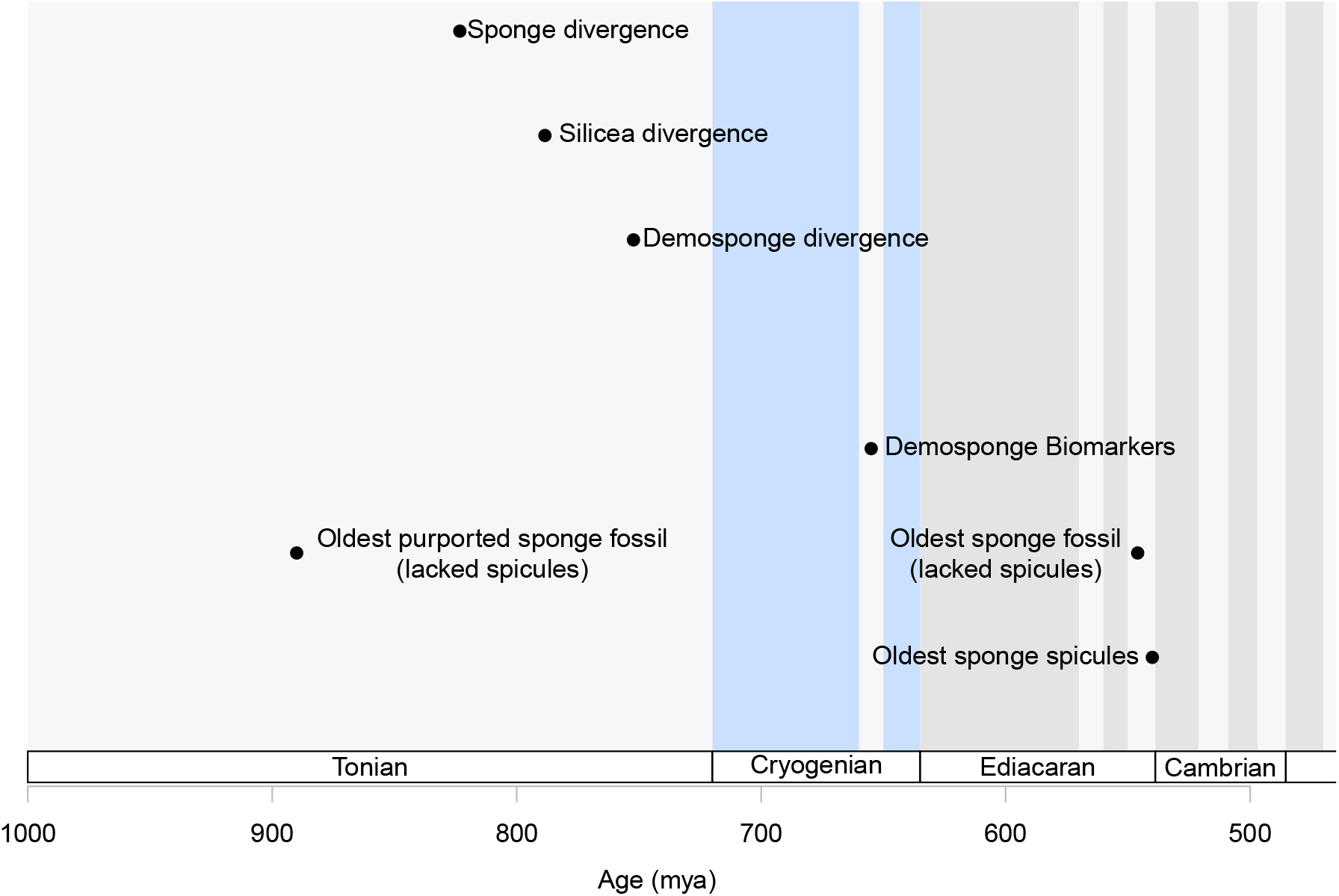
Conflicting fossil, molecular clock, and biomarker evidence for the origin and early history of siliceous sponges. A missing spicule record of at least 100 million years is implied by span of time before the oldest spicule fossil in the late Ediacaran and the presence of Cryogenian aged lipid biomarkers. The missing record is worse when considering the Cryogenian or Tonian molecular clock divergence times for siliceous sponge clades (dates from 1, 6, 11, 26, and 38). Cryogenian glaciations are denoted in vertical blue bars and subdivisions of geological time by grey bars.

At the most extreme, if spicules evolved soon after the divergence of hexactinellids and demosponges, then there is at least 100 million years between the appearance of spicules in the latest Ediacaran. For a spicule gap of any duration to occur, taphonomic processes would have to produce the wholesale exclusion of spicules from the rock record. Alternatively, if spicules evolved late in each group’s history, then there need not be a taphonomic spicule gap at all. A third potential alternative is that the deep history implied by the biomarker and molecular phylogenetic records are both wrong (10, 17-19) and instead sponges evolved and radiated over a compressed timeline in line with their appearance in the body fossil record. Despite the general uncertainties in reconciling molecular clock divergence time estimates and the fossil record (18-21), we suspect this third hypothesis is unlikely because bilaterian body fossils are present in Ediacaran faunas (6, 22) coincident with the oldest sponge body fossils (11, 16, 23) and just prior to the oldest preserved spicules (24-26).

To produce a spicule gap lasting over 100 million years would have required a general and consistently operating global mechanism. High sea surface temperatures are likely to be the only viable candidate due to their global extent and because high temperatures increase the kinetics of diagenetic pathways that impact biogenic silica (27-29) and could be the source of high levels of dissolved silica levels in the Ediacaran (30-32). Siliceous diatom frustules (e.g., biogenic Si, bSi, or opal-A) are known to dissolve if exposed to high temperatures and in undersaturated conditions (e.g., 33, 34). In fact, this susceptibility to temperature driven dissolution may be the cause of a paucity of fossils during the Mesozoic until global temps cooled sufficiently to permit preservation (35). Given the extreme heat of the Ediacaran and Cambrian, high temperature-driven dissolution this process may also have the potential to cause the Ediacaran sponge spicule gap.

Assessing the pattern of paleotemperature across this interval is complicated by the fact that the best isotopic estimates of paleotemperature are measured in shelly fossils, which did not originate until the late Ediacaran and Early Cambrian and then only became abundant in the Cambrian. Clumped isotope paleothermometry provides a means of estimating paleotemperatures from carbonate rocks that predate the origin of calcitic skeletons and permit the direct comparison of temperatures spanning the Ediacaran and Cambrian (36). Although the precision of clumped isotope paleothermometry is likely to be affected by diagenetic alteration, it remains the only method to compare relative changes in paleotemperature prior to the origin of carbonate skeletons.

To assess whether there is an absence of evidence or evidence of absence of spicules, we compiled a comprehensive spicule and body fossil dataset for late Ediacaran and Cambrian siliceous spicule bearing sponges derived from the literature (Paleobiology Database, Geobiodiversity Database, and undatabased monographs, see Supplemental Information) and global digitized museum collections (derived from the Global Biodiversity Information Facility). This dataset contains every known siliceous sponge occurrence, but lacks undigitized museum samples and spicules present in rock samples housed in museum collections but not sampled for spicules. Combined with a paleothermometer that spans the Ediacaran and Cambrian (36), this dataset allows us to assess the likelihood of a taphonomic spicule gap. We can distinguish between absence of evidence and evidence of absence if spicule preservation occurs in the Cambrian at paleotempeatures higher than inferred in the Ediacaran. If this is observed, then we can infer that spicules should be preserved if they in fact were present during the Ediacaran.

## Materials and Methods

Fossil sponge spicules are preserved in association with body fossils or as isolated spicules. Sponge species tend to possess many shapes and types of spicules and so it is often not possible to link an isolated spicule to a species, genus, or family although it is possible to identify higher taxonomic association, e.g., hexactinellids and demosponges. For this reason sponge fossils in the broad sense occupy two distinct modes, the body fossil record with low-level taxonomic IDs that inform evolutionary work, and the isolated spicule record, that highlights only the presence and abundance of higher-level clades of sponges. For our question concerning the timing of the evolution of spicules and if there is a large-scale taphonomic process that could preferentially removed spicules from the fossil record, we can use both the body and spicule records.

No single resource possesses all information about Ediacaran and Cambrian sponge samples. Therefore we complied a comprehensive dataset consisting of all known spicules and body fossil occurrences. There are surely spicules we have not sampled to be found in undigitized museum drawers or in unstudied rock samples. We assembled a dataset that derives and pools occurrences form multiple data sources that covers all known published and digitized Ediacaran and Cambrian sponge spicules and body fossils.

To illustrate why we need to combine data sources, we walk through the limitations of each data source individually. This is not to highlight flaws in those data sources, but to instead focus on the scientific opportunities we can make when combining data from multiple resources into more comprehensive datasets. Specimens in natural history museum collections represent a vast under tapped source of information (37) for large scale analyses, with the majority unpublished in the peer-reviewed literature, not described in formal taxonomic works and therefore often cataloged with limited taxonomic resolution, or not just digitized and made publicly available. Because we care only about high-level taxonomic identity for this work, we do not have issues with limited taxonomic resolution of digitized occurrences from museum collections. Another issue that arises is that the full paleontological literature may not be entered into global fossils databases or that these databases may systematically lack literature that is not published in English and therefore introduce subtle (or not so subtle) distortions in spatial sampling. For example, large-scale datasets such as the Paleobiology Database (paleobiodb.org) primarily rely on species or genera identified from body fossils in the peer-reviewed published literature. This is because the main mission of the database has been to generate a standard datum, the occurrence, that can be used for global scale sample standardization (38, 39). Monographs describing isolated spicule samples may be excluded on purpose to maintain data quality or for convenience in data entry. This does not necessarily mean that isolated spicules are universally excluded from the Paleobiology Database, but instead that there is no systematic sampling of the literature describing isolated spicules the way that there is for body fossils. The Geobiodiversity Database (gbdb.org run out of the Nanjing Institute of Paleontology (40-43) compliments the Paleobiology Database by adding fossil coverage especially in Ediacaran and Cambrian occurrences in China, and adds new data in the amount and type of non-fossiliferous sedimentary rocks in an interval. The Global Biodiversity Information Facility (GBIF.org) hosts a wide range of digitized museum collection occurrences including material that has not been published in peer-reviewed literature. Depending on circumstances and resources, museum collections can lack updated taxonomy and have outdated chronostratigraphic information, but they routinely record precise geolocation data for fossil samples and maintain strong stratigraphic and locality records. Additional information, not captured by the main databases, is recorded in the paleontological literature not yet entered into the Paleobiology Database or GBDB or with sampling that does not conform to their needs and requirements.

Another issue that makes combining data from these multiple databases complicated is that each utilizes its own set of fields, conventions, and standards. For us, the primary occurrence datum consists of a unique stratigraphic collection locality that possesses the presences of a sponge fossil (isolated spicule or body fossil). The Paleobiology Database, GBDB, and GBIF all share occurrences, have chronostratigraphic, geographic, and potentially taxonomic information, but in particular for GBIF, the important chronostratigraphic and taxonomic information can be spread across multiple fields and so requires some data cleaning and curation to integrate easily with data from the other data providers.

Paleobiology Database: Our dataset from the Paleobiology Database is derived from two downloads (made 10/1/24): One explicitly downloading occurrence data on Ediacaran and Cambrian sponges using their API version 2.1: https://paleobiodb.org/data1.2/occs/list.csv?datainfo&rowcount&base_name=Porifera&interval=Ediacaran,Cambrian&pgm=gplates&show=full and one pulling all Ediacaran and Cambrian occurrences regard-less of phylum level taxonomic IDs: https://paleobiodb.org/data1.2/occs/list.csv?datainfo&rowcount&interval=Ediacaran,Cambrian&pgm=gplates&show=full. Archeocyathids and calcitic sponges were excluded after download so the final dataset includes only hexactinellid, demosponge, and isolated siliceous spicules.

Geobiodiversity Database: We downloaded the complete dataset on 6/18/24 and culled the dataset to include only occurrences of Ediacaran and Cambrian hexacinellids, demosponges, and isolated spicules. We standardized timescale dates using regional timescale definitions (45-47). Global Biological Information Facility: The GBIF data was generated by searching Basis of Record = Fossil Specimen; Scientific Name = Porifera which yielded 65,812 occurrence records. A Darwin Core (DwC) Archive was then downloaded to get the full suite of data including geologic context information. Our 1/8/25 GBIF Occurrence Download doi is: https://doi.org/10.15468/dl.s7f3un. Next we filtered out records younger than Cambrian by examining the multiple chronostratgraphic fields present in downloaded occurrence dataset file. We then reviewed records that have no geologic time reported, deleting occurrences without lithostratigraphy and otherwise dating to stage/epoch level when sufficient lithostratigraphic data was present to determine in reverse the chronostratigraphic context. Records lacking geologic context data and were removed. We removed occurrences of Calcarea, archaeocyathids, 37 cephalopod records that erroneously had the higher taxonomy listed as porifera, and *Brooksella* records including synonyms (e.g., *Laotira cambria*). Examination of fields with information related to preparations revealed additional information that occurrences represented archaeocyathid thin section or other determination information that allowed them to be identified as not relevant and excluded from the dataset. Final result is 1557 occurrence records.

Correlating across regional timescales can be difficult in the Cambrian (48), and so translating occurrences resolved to intervals in one region to a global timescale can mask temporal resolution. Instead of correlating each occurrence to a global timescale, we used the published chronostratigraphic data associated with the occurrence record, thus utilizing regional timescales and their associated temporal intervals when provided. Using regional timescales (defined in 44–47) does not inflate occurrences because the chronostratigraphy of each occurrence can only be denied once.

To asses mode of preservation, we reviewed references for occurrences to extract descriptions pertaining to the preservation of spicules in the fossil record. This occurrence data was accessed and downloaded from the Paleobiology Database (www.paleobiodb.org) on 10/1/24, with base name as Porifera and time interval as Cambrian. The spreadsheet included a total number of 1365 occurrences of 155 Cambrian sponge species from 78 references. Occurrences of Porifera in the classes Archaeocyatha and Calcarea were removed from the dataset. Subsequently, these descriptions were categorized as OS [original silica], OR [original silica with partial replacement, e.g., limonite-stained or coating], R [replacement], CMI [cast, mold, and impression] (see Table 1 for verbatim definitions).

**Table 1:**
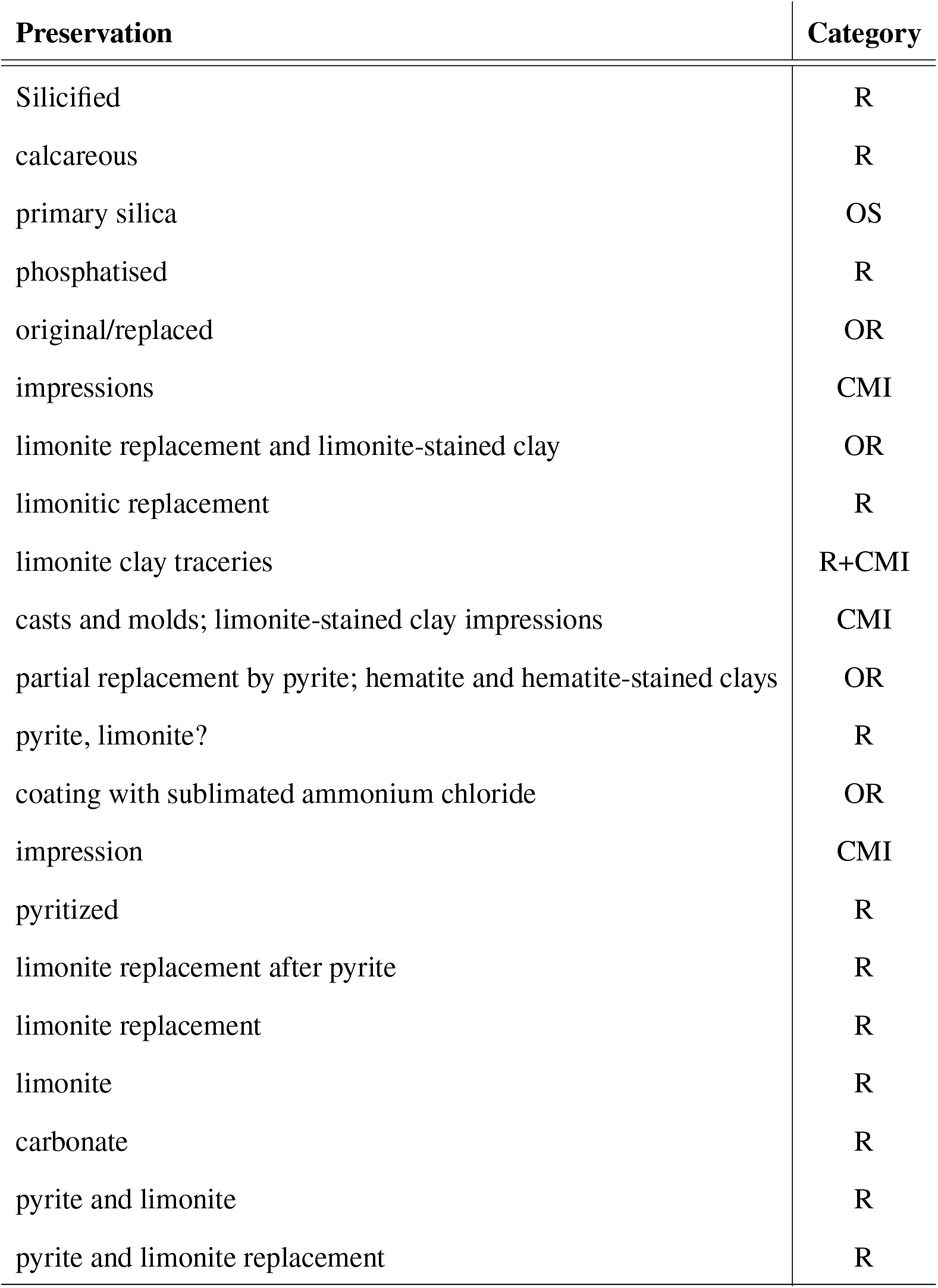
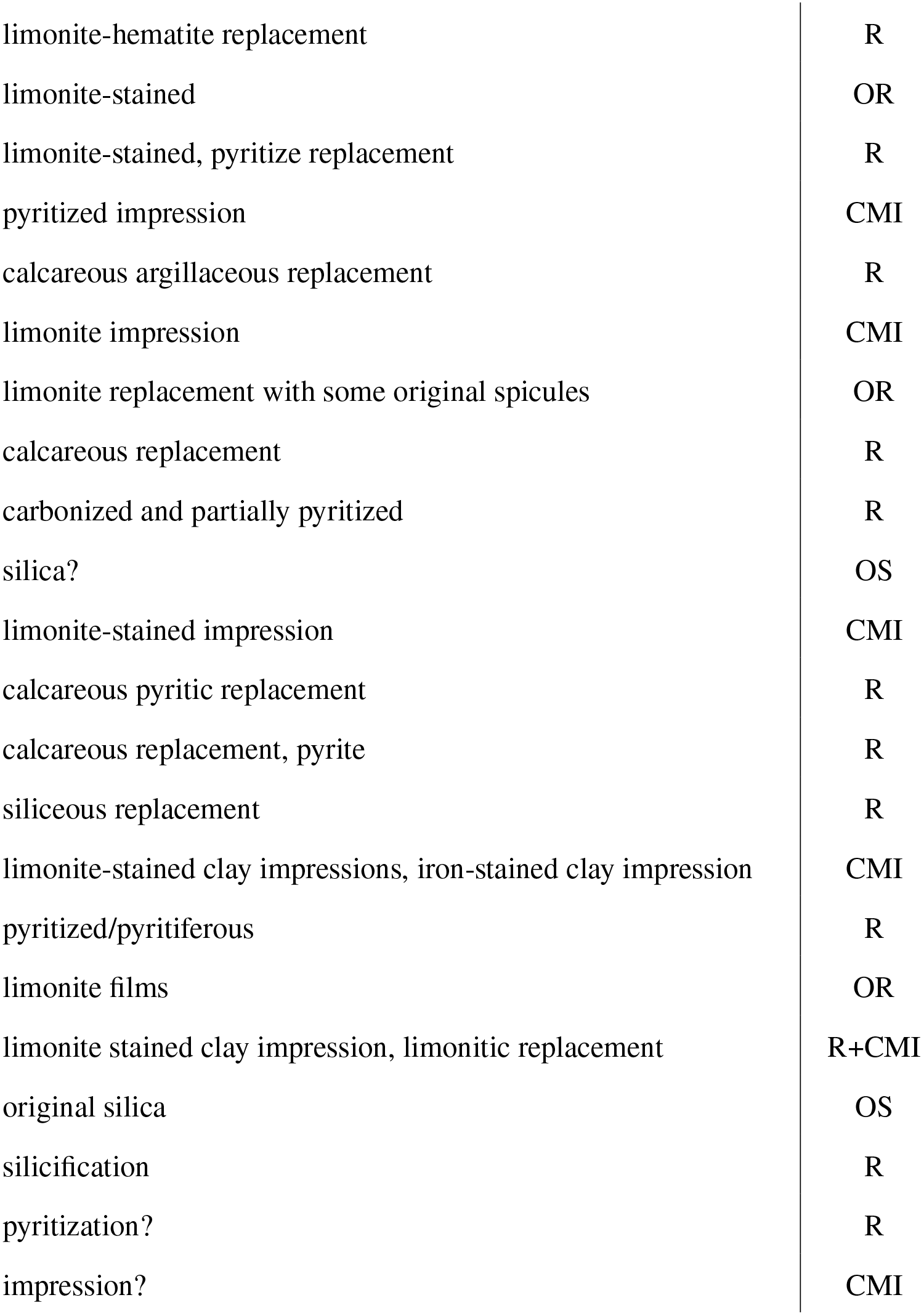
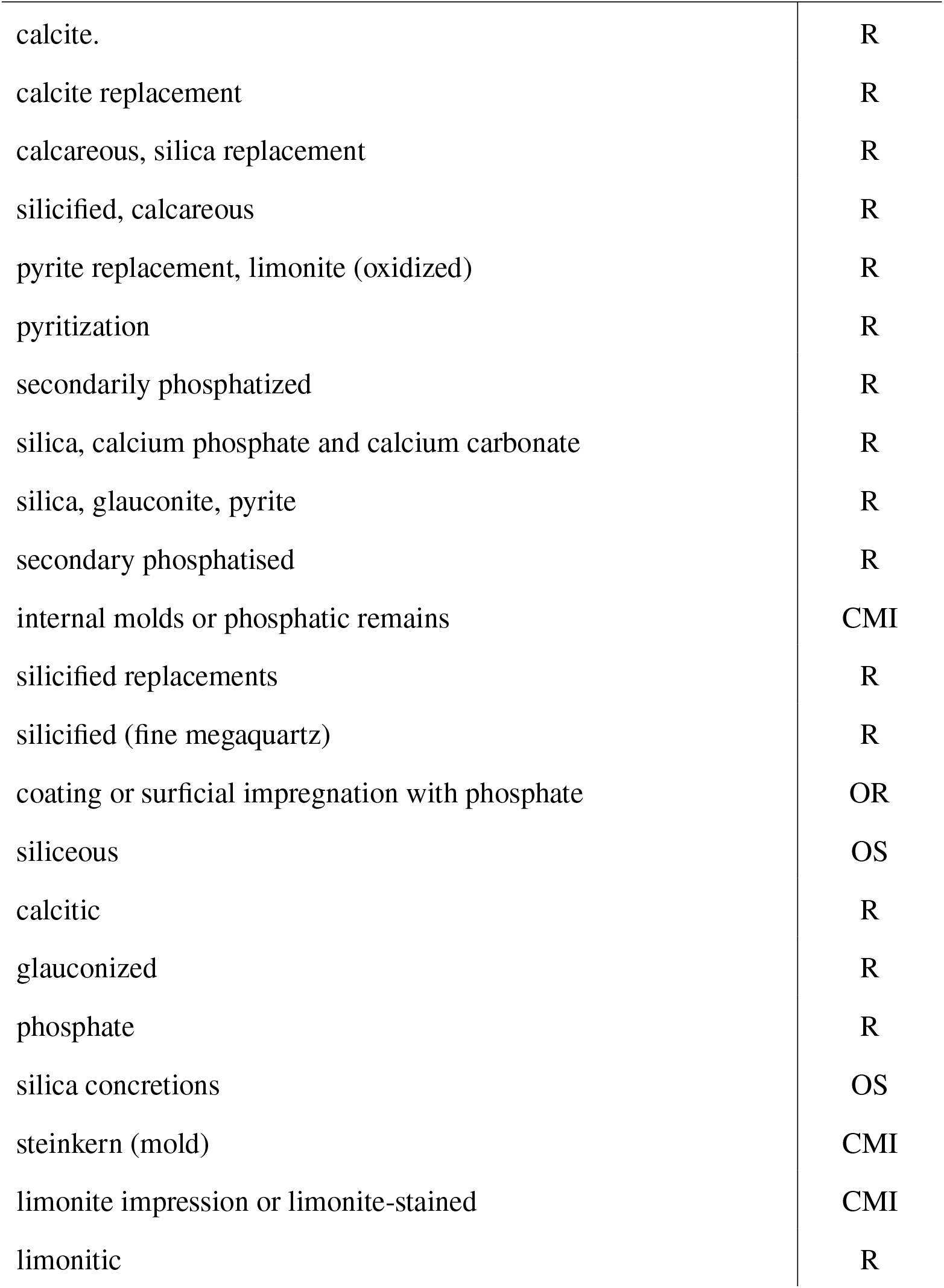
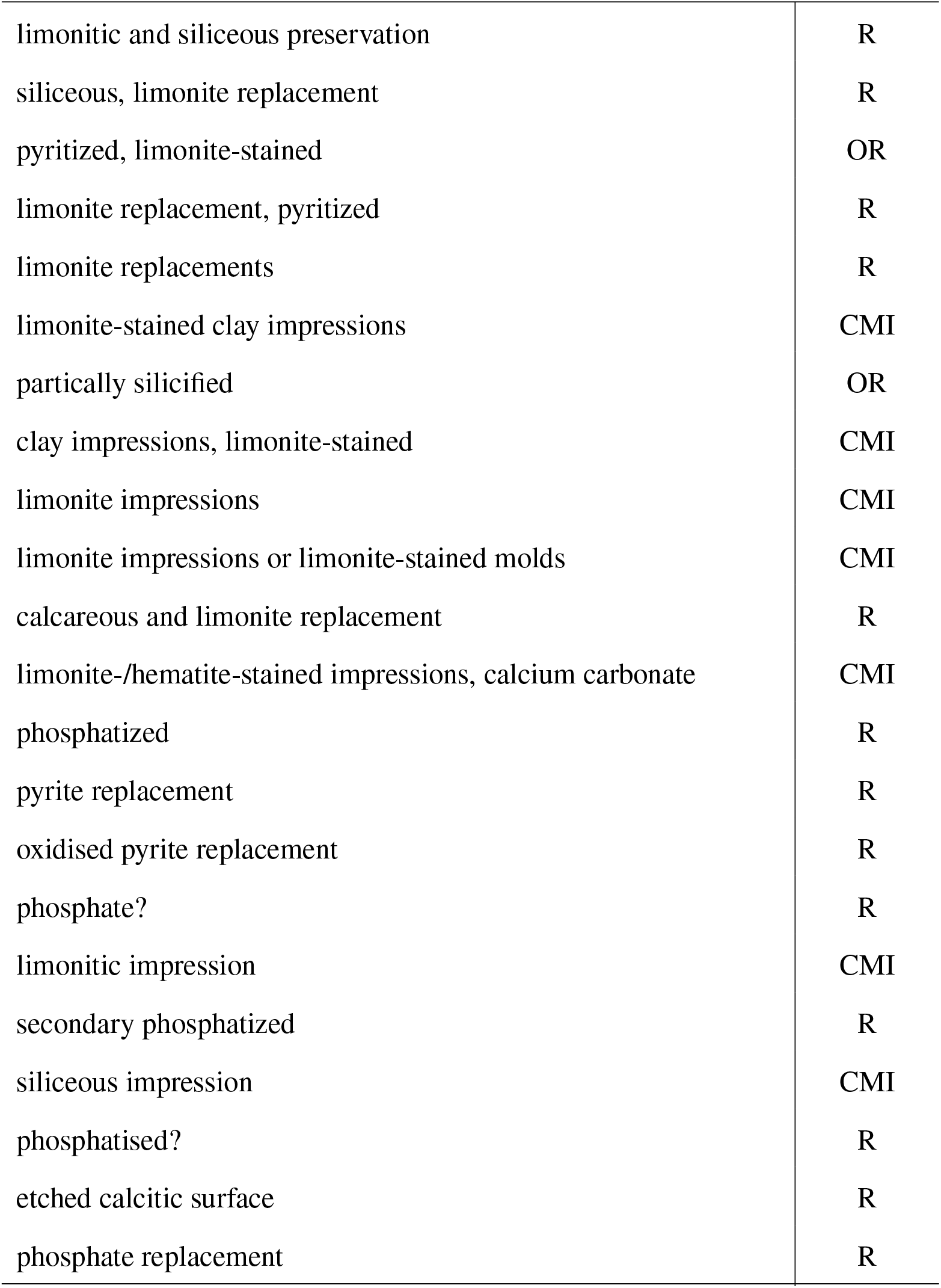
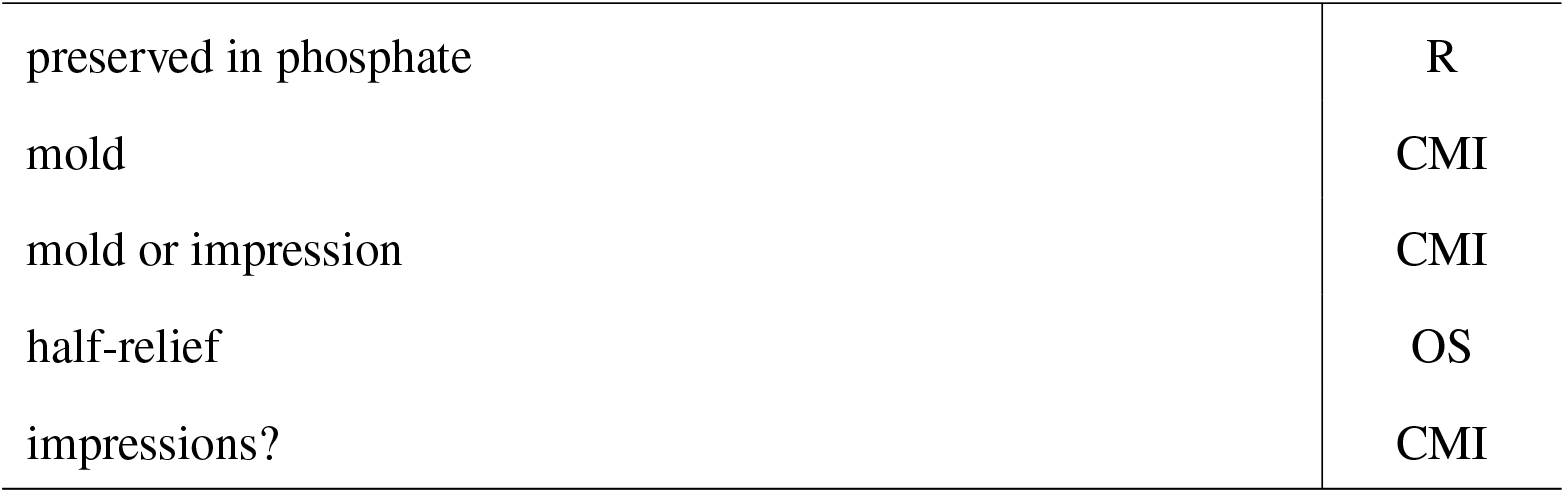
Definitions of modes of preservation.

## Results

The results are strikingly clear: siliceous sponge spicules are routinely preserved in the Cambrian (Fig. 2) during intervals where temperature is inferred to far exceed those observed in the Ediacaran (Fig. 3). These observations are consistent with the low solubility of sponge spicules even at high experimental temperatures (29, 32, 33). Cambrian intervals with spicule occurrence numbers on par with Ediacaran occurrences include some of the hottest intervals of the Cambrian, and on average would likely have exceeded temperatures of more than three quarters of the Ediacaran. Many dozens of occurrences are known from the hottest Cambrian time interval, that would have matched even the heat that followed the Marinoan glaciation. The few late Ediacaran spicule occurrences occur at temperatures far below those observed for all Cambrian Occurrences (Fig. 4). Moreover, even though fossil sampling may also be controlled by the amount of rock available to sample (37), Cambrian rock volumes only exceed Ediacaran levels in the middle Cambrian (46), 20 million years after the sponge spicule record begins.

**Figure 2.**
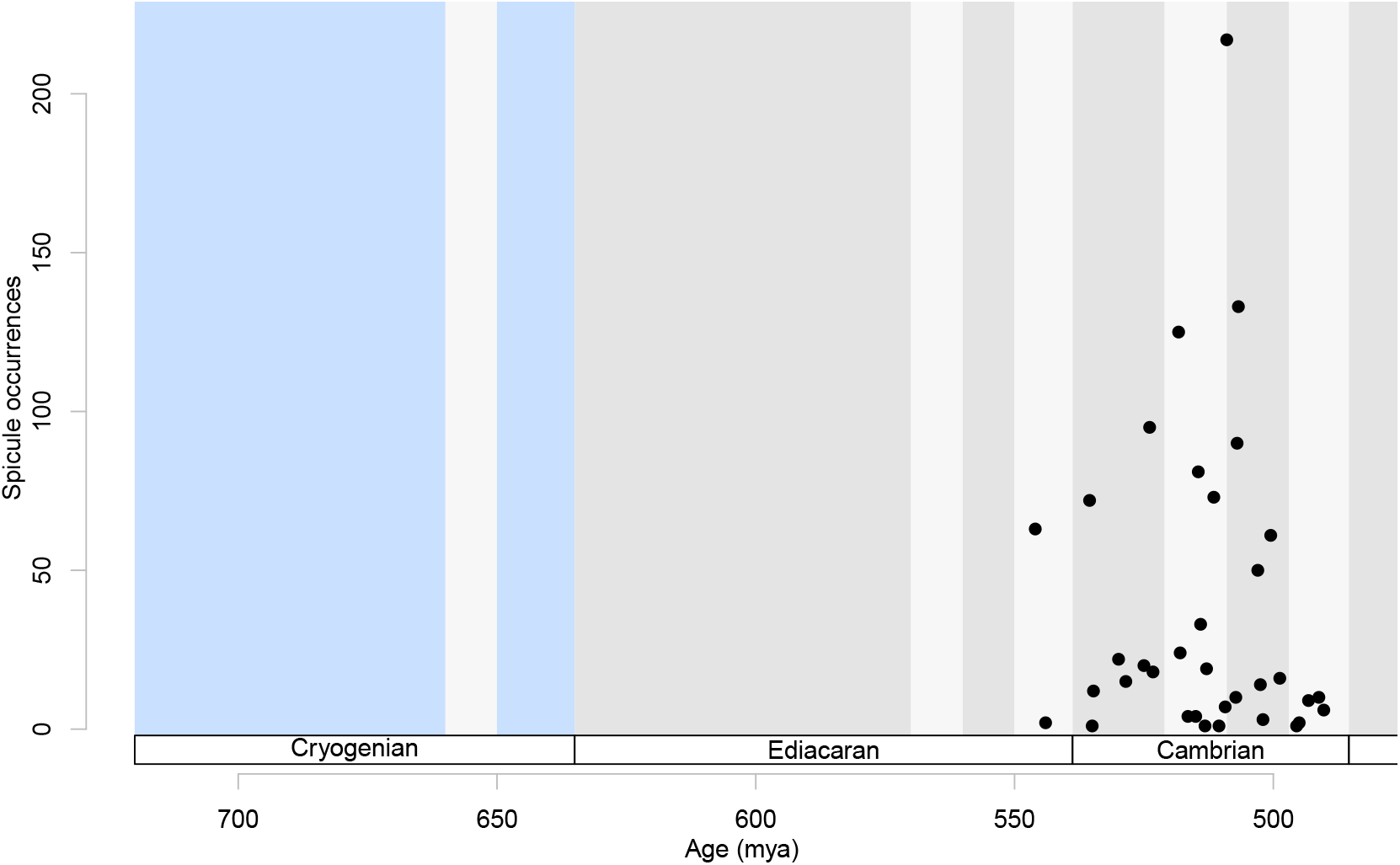
Occurrences of silicious sponge fossils in the Ediacaran and Cambrian. Data are derived from a comprehensive dataset of spicule and body fossils derived from the literature (PBDB, GBDB, and undatabased monographs and literature) and digitized museum collections data (derived from GBIF). Each fossil occurrence records the presence of spicules or body fossils at a unique geological time and locality. Occurrences dateable to only regional timescales were left uncorrelated with the global timescale and plotted separately. Occurrences are dates to epoch or stage level are plotted in the center of those intervals. Ediacaran occurrences are known to occur at the very top of the sections but are here shown to be slightly older because of the coarseness of the Ediacaran timescale. Blue vertical bars denote the Cryogenian “Snowball Earth” glaciations.

**Figure 3.**
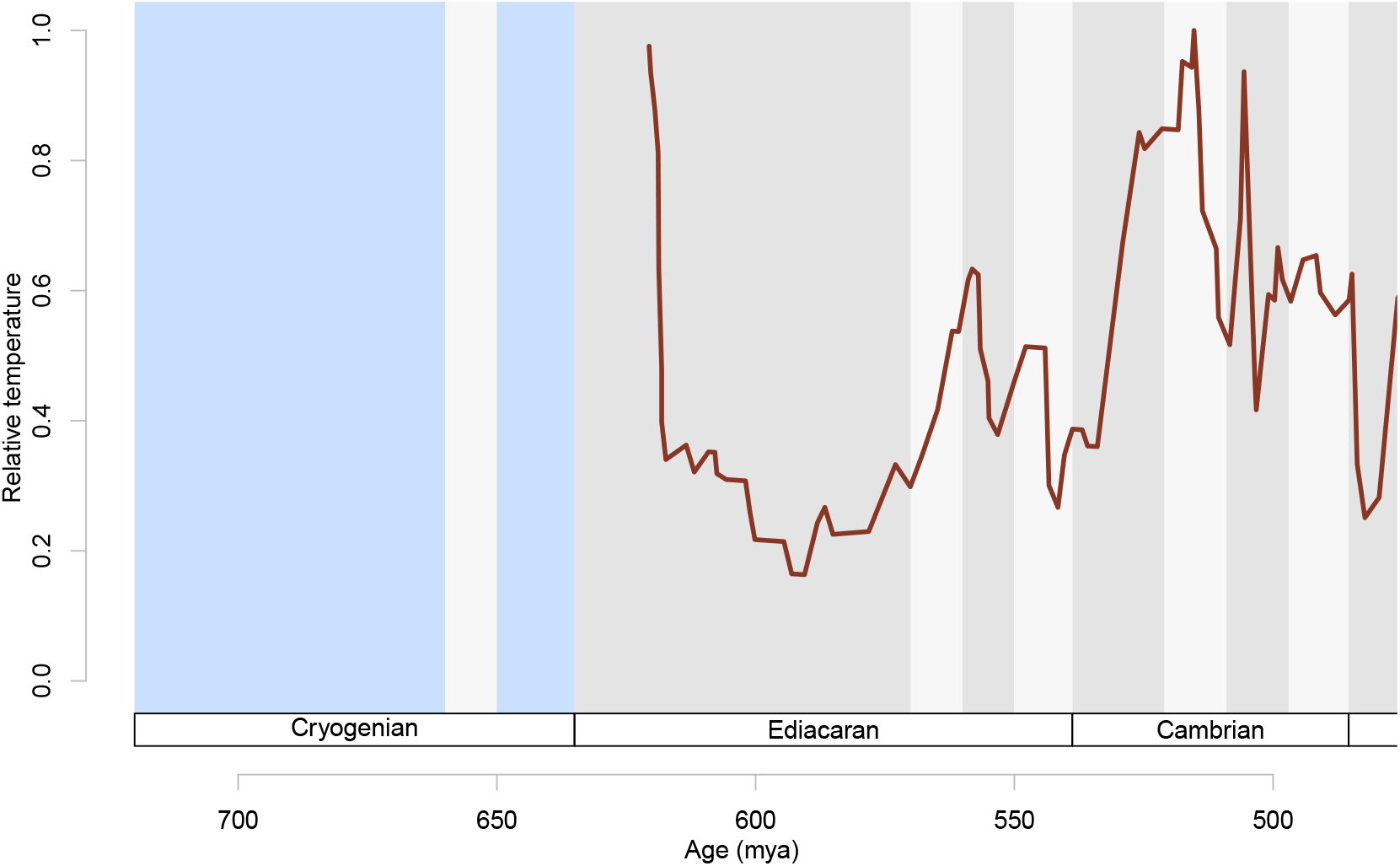
Pattern of global sea surface temperatures for the Ediacaran and Cambrian. Data are derived from reference 35, a compilation of clumped isotopic data. Because temperatures derived from clumped isotopes tend to run hot due to diagenetic effects, we show minimum clumped temperature data and rescale the temperatures to the maximum temperature in this temporal span to show relative changes only.

**Figure 4.**
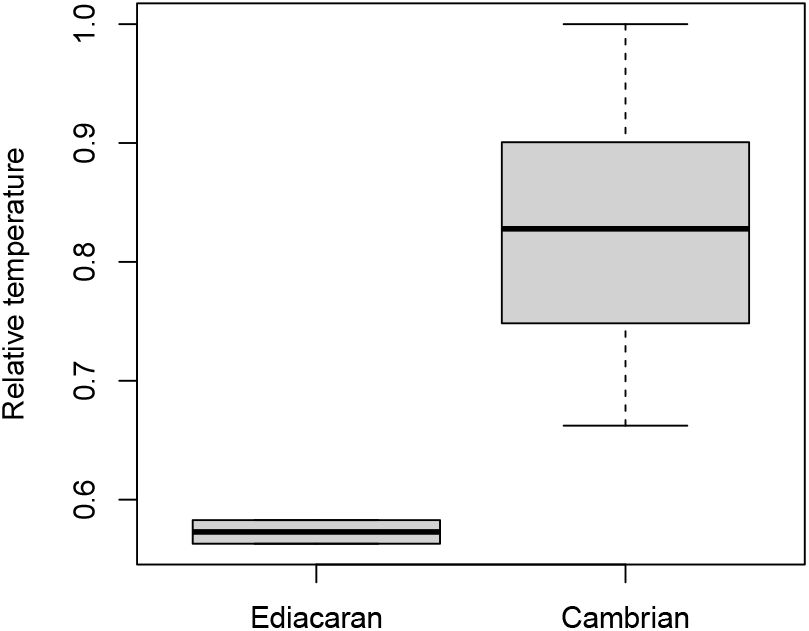
Relationship between temperature and spicule preservation. For intervals with spicule occurrences, Cambrian paleotemperatures far exceed those inferred for the Ediacaran (t-test, t = 12.065, p < 0.001).

If present, spicules would have been more likely to have been preserved in the Ediacaran than the Cambrian for two reasons. First, because of the cooler temperatures, biogenic silica (bSi) reaction kinetics should be slower than when temperatures are higher (33, 34). And second, because the high dissolved silica concentrations ([dSi]) in Ediacaran oceans (31, 32) would make spicule dissolution more difficult due to the shallower concentration gradient than would occur in a silica poor ocean. Both processes limit the ability of silicious spicules to dissolve or be otherwise lost prior to burial and lithification of sediments. Nevertheless, diagenetic processes can occur post-burial and post-lithification. To estimate the frequency of post-burial loss we tabulated of modes of preservation using a subset of total occurrences (see supplementary data) based on assessments of modes of preservation described in the primary literature (assessment of preservation mode of museum specimens is not feasible with currently available data, excluding them from this dataset/analysis). We find that in the Cambrian, mineral replacements, molds, and casts of siliceous spicules make up the bulk of the record (∼86%, Table 2, Fig. 5).

**Table 2:** Number of occurrences for each type of preservation in the Paleobiology Database subdataset. Preservation types include OS (original silica), OR (original silica with partial replacement), R (replacement), and C (cast, mold, and impression.

| Preservation Types | Original silica remains |  | Silica replaced or absent |  |  | Total |
| --- | --- | --- | --- | --- | --- | --- |
|  | **OS** | **OR** | **R** | **CMI** | **R+CMI** |  |
| Occurrences | 17 | 39 | 272 | 78 | 4 | 410 |

**Figure 5.**
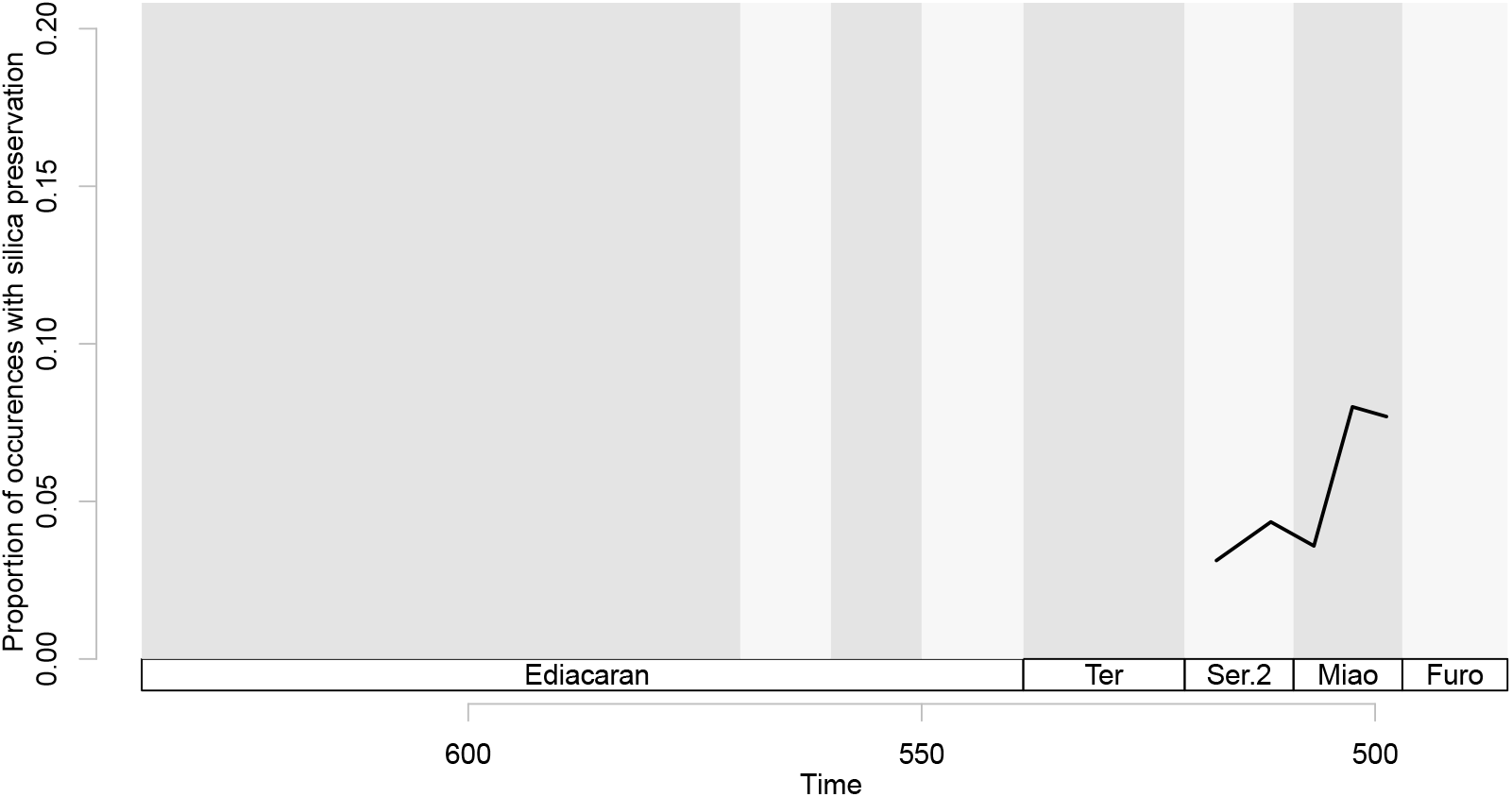
Silica preservation in Cambrian sponge spicules. Preservation mode was assessed for Cambrian siliceous spicules. The majority of Cambrian spicules are not preserved as original silica. Here we show the proportion of spicules with silica preservation.

## Discussion and Conclusions

From this evidence, we infer that siliceous spicules were absent during the bulk of the Ediacaran. If spiculate sponges were present in the Ediacaran, we would expect to find preserved spicules even if only as casts, molds, or replaced spicules and identifiable as such.

It is not until the latest Ediacaran that identifiable spicules appear in the rock record (24-26). Prior to that, evidence is emerging that sponges were present but lacked spicules. For example, *Helicolocellus cantori*, a stem hexactinillid lacking spicules, occurs in the late Ediacaran (11). Fossils possessing patterns of pores diagnostic of a sponge pump, such as *Thectardis* are known from Ediacaran Mistaken Point faunas of Newfoundland (16). Purported aspiculate keratosin sponge fossils have also been described from the Tonian Little Dal formation in Canada (39).

At first glance, the hypothesis of a taphonomically-caused spicule gap seemed to be plausible, but key biogeochemical and phylogenetic assumptions have recently been revised. Historically, sponge spicules were thought to be a negligible sink of biogenic silica, playing little role in the global marine silica cycle budget summaries (40-42). But sponge spicules, unlike some diatom frustules (42, 43), are extremely resistant to dissolution (45-48) even in more extreme temperatures (85°C) and alkaline leachate concentrations (46, 49) than would have occurred in the Ediacaran and Cambrian. The second assumption that maintained the plausibility of the taphonomic spicule gap hypothesis is that siliceous spicules evolved once in the history of silicious sponges and are homologous in hexactinellids and demosponges (50-54). Although differences in how spicules form during development have long been identified (55, 56), recent molecular phylogenetic analyses support a sister relationship between hexactinellids and demosponges (8, 9, 14, 50, 57) and therefore the most parsimonious interpretation is a homologous origin of silicious spicules. Yet, the long-held assumption that siliceous spicules evolved only once is now challenged by a new understanding of the mechanisms of spicule formation (58, 59). In demosponges, silicateins – enzymes homologous to cathepsin L proteases – catalyze the polycondensation of silicic acid to form biosilica, initiating axial growth and forming the central filament of spicules and support appositional growth by promoting lateral silica deposition (60-65). In contrast, hexactinellid sponges lack silicateins entirely and instead employ a suite of unrelated proteins – hexaxilin, glassin, and perisilin – to direct biosilica deposition. Among these, hexaxilin is the most broadly distributed, with homologs found in all examined hexactinellid taxa, and it serves as the axial filament around which silica is deposited (12, 59), whereas glassin and persilin are less widely distributed but are involved in spicule thickening (59, 66, 67).

Outside of either demosponges or hexactinellids, homoscleromorph sponges are sister to calcareous sponges and appear to reflect an additional case of independent spicule evolution. Homoscleromorphs have siliceous spicules but lack silicateins (68), and uniquely, spicule formation occurs in epithelial cells rather than in dedicated sclerocytes (58). Moreover, independent origins of other spicule biominerals, such as calcium carbonate in calcarious sponges (69), potentially many lineages of hypercalcifying sponges (70) and hypercalcifying demosponges (which possess silicious spicules) highlight the commonality of multiple origins of biomineralization within sponges. Together, these molecular and developmental differences strongly support the hypothesis that siliceous spicules evolved independently at least three times in sponges.

These recent discoveries together with our results lend further credence to the interpretation that sponges produced the sterane 24-isopropylcholestane found in Cryogenian interglacial aged sediments in Oman (1, 3, 5, 59) because sponges have a demonstrable pre-spiculate evolutionary history. All together, we think that the evidence points to a deep history of the hexactinellid and demosponge clade prior to their convergent evolution of siliceous spicules in the late Ediacaran and early Cambrian. Consequently, we can not only take the fossil record at face value, but we may also be able to take the molecular divergence time estimates and the lipid biomarkers at face value.

Consequently, silicious sponges join the ranks of other biomineralizing animal groups (71, 72) and microorganisms in that they evolve biomineralization at the Ediacaran – Cambrian boundary (with the possible exception of bryozoans (73, 74)). The evolution of biomineralization may relate to the influx of weathering products into the ocean during and up to the end of the Great Unconformity (75), or through some other, yet unknown, mechanism. Whatever the ultimate cause, the number of independent origins of all forms of biomineralization and the Cambrian radiation of animal groups at this time are likely to be more than synchronicity. The question now is whether the origin of biomineralization drove the radiation of animal groups or whether biomineralization and animal diversification had a common cause.

## Acknowledgments

We thank Karen Chin, Melanie Hopkins, Sarah Leventhal, Jack Shaw, and Lizzy Trower for helpful discussions. Funding was provided by National Science Foundation Award EAR-2207109 (CS) and National Science Foundation Award GEO CI-2324688 (TSK, CS).

